# TWAS coupled with eQTL analysis reveals the genetic connection between gene expression and flowering time in Arabidopsis

**DOI:** 10.1101/2022.12.07.519424

**Authors:** Pei-Shan Chien, Pin-Hua Chen, Cheng-Ruei Lee, Tzyy-Jen Chiou

**Affiliations:** Agricultural Biotechnology Research Center, Academia Sinica, Taipei, Taiwan; Institute of Ecology and Evolutionary Biology, National Taiwan University, Taipei, Taiwan; Institute of Plant Biology, National Taiwan University, Taipei, Taiwan

**Keywords:** *Arabidopsis thaliana*, genome-wide association, transcriptome-wide association, natural variation, flowering time, haplogroup, expression quantitative trait locus mapping

## Abstract

Genome-wide association study (GWAS) has improved our understanding of complex traits, but challenges exist in distinguishing causation versus association caused by linkage disequilibrium. Instead, the transcriptome-wide association study (TWAS) detects direct associations between expression levels and phenotypic variations, providing an opportunity to better prioritize candidate genes. To assess the feasibility of TWAS, we investigated the association among transcriptomes, genomes, and various traits, including flowering time in *Arabidopsis*. First, the associated genes formerly known to regulate growth allometry or metabolite production were identified by TWAS. Then, for flowering time, six TWAS-newly identified genes were functionally validated. Analysis of expression quantitative trait locus (eQTL) further revealed a *trans*-regulatory hotspot affecting the expression of several TWAS-identified genes. The hotspot covers the *FRIGIDA* (*FRI*) gene body, which possesses multiple haplotypes differentially affecting the expression of downstream genes, such as *FLOWERING LOCUS C* (*FLC*) and *SUPPRESSOR OF OVEREXPRESSION OF CO 1* (*SOC1*). We also revealed multiple independent paths towards the loss of *FRI* function in natural accessions. Altogether, this study demonstrates the potential of combining TWAS with eQTL analysis to identify important regulatory modules of the *FRI-FLC-SOC1* for quantitative traits in natural populations.

**Highlight:** Combining TWAS with eQTL analyses identifies haplotypes connecting flowering genes with their physiological trait, strengthening the importance of *FRI-FLC-SOC1* regulatory module on flowering time among the Arabidopsis natural population.

## Introduction

Genome-wide association studies (GWAS) have brought about remarkable advances in dissecting complex genetic traits and identifying associations between single nucleotide polymorphism (SNP) markers and terminal phenotypes (Atwell *et al*., 2010). Potential causal genes can then be identified in the regions with high linkage disequilibrium (LD) with candidate SNPs. However, it remains challenging to translate from statistical associations to molecular causations. Considering that LD distance can range from 1 kb in maize (*Zea mays*; Tenaillon *et al*., 2001) and 10 kb in Arabidopsis (*Arabidopsis thaliana*; Kim *et al*., 2007) to an average of 100 kb in soybean (*Glycine max*; Zhou *et al*., 2015), the number of candidate genes can range from few to many. Although GWAS can quickly and efficiently delineate genome regions that control traits and provide markers to accelerate breeding by marker-assisted selection, most variants reside outside protein-coding regions, resulting in difficulty understanding the biological mechanism underlying these revealed associations.

To circumvent the difficulty and to better prioritize causal genes, one possible solution is to illuminate the genetic information that connects to traits through multiple intermediate processes, including transcription, translation, epigenetic modification, and metabolism (Kremling *et al*., 2019). For example, using simple linear regression, the associative transcriptomics of traits was proposed to identify genes related to oil quality in the polyploid rapeseed (*Brassica napus*) (Harper *et al*., 2012; Havlickova *et al*., 2018). A model-based transcriptome-wide association study (TWAS) was also developed to improve the power of gene discovery recently. In maize, a TWAS method termed expression read depth GWAS (eRD-GWAS) that employs a Bayesian-based model was developed (Lin *et al*., 2017). A flowering time-associated candidate gene, *MADS TRANSCRIPTION FACTOR69* (*ZmMADS69*), has since been identified and functionally characterized (Liang *et al*., 2019). In addition, eRD-GWAS identified root trait-associated candidate genes previously known to affect root architecture (Zheng *et al*., 2020). In rapeseed, Tang *et al*. (2020) combined two sets of transcriptome data from developing seeds sampled on different days after flowering for TWAS using a mixed linear model. A *BnPMT6* gene was thus revealed to regulate seed oil content negatively for the first time (Tang *et al*., 2020). These reports support the potential value and feasibility of TWAS in plants.

Recent studies that combined GWAS, TWAS, and expression quantitative trait locus (eQTL, identifying genetic variants affecting the expression of specific genes) uncovered the genetic regulatory network that orchestrates the initiation of fiber development (Li *et al*., 2020) and male sterility (Ma *et al*., 2021) in cotton. Due to the importance of flowering, an essential process for developmental transition and a crucial trait for crop yield (Blumel *et al*., 2015), the mechanism of controlling flowering time has been well characterized (Bouché *et al*., 2016). Studies in Arabidopsis have identified individual signaling pathways and gene components within these pathways that affect flowering time in different plant species, deciphering the complexity of this regulatory network (Kim, 2020). Also, genetic analyses of Arabidopsis revealed how the activities of these pathways are altered in nature and how the effects of environmental stimuli on flowering time are balanced to adapt to different geographical locations (Caicedo *et al*., 2004; Fransz *et al*., 2016). By taking advantage of the publicly available dataset of Arabidopsis, we analyzed transcriptomes, genomes, and several traits, including flowering time, from 1,001 Genomes to examine the feasibility and capability of TWAS using mixed linear model (MLM), a model commonly implemented in the conventional GWAS.

In this study, we first provided evidence to show the potential of TWAS in identifying associated genes related to growth allometry, metabolite production, and flowering time in Arabidopsis. Then, we functionally validated TWAS-identified candidate genes for flowering time. We further integrated eQTL analysis into the TWAS result and identified a *trans*-regulatory hotspot that covered the *FRIGIDA* (*FRI*) gene body on the chromosome (Chr) 4. Accordingly, we found that the *FRI* haplotype connects transcript levels of its own and downstream regulators, *FLOWERING LOCUS C* (*FLC*) and *SUPPRESSOR OF OVEREXPRESSION OF CO 1* (*SOC1*) to the terminal phenotype, flowering time. Through coupling with eQTL analysis, the genetic association between causal genes and physiological phenotypes was revealed, shedding light on the importance of the *FRI*-*FLC*-*SOC1* signaling module in regulating flowering time from the genetic point of view.

## Materials and methods

### Transcriptome data

RNA-seq-derived transcriptomic data from rosette leaves of 727 Arabidopsis (*Arabidopsis thaliana*) accessions grown under ambient temperature in greenhouses from a previous study (Kawakatsu *et al*., 2016) were downloaded (http://neomorph.salk.edu/1001.php). After trimming by Trimmomatic (Bolger *et al*., 2014), reads were mapped using HISAT2 (Kim *et al*., 2015) to the Araport11 (Cheng *et al*., 2017) genome and annotation. Next, the expression of the annotated genes was quantified and batch normalized by StringTie (Kovaka *et al*., 2019), resulting in transcripts per million (TPM) as expressional values for each gene. Finally, chloroplast and mitochondria genes were removed, retaining 36,631 genes in each accession inbreed line.

### TWAS

Various phenotypic datasets of Arabidopsis were downloaded from AraPheno (https://arapheno.1001genomes.org/). For TWAS analysis, we removed genes with zero or nearly zero variance (> 95% of the lines sharing the same transcript level) across lines using nearZeroVar function from the R caret package. Thus, the number of explanatory variables associated with individual traits may differ (i.e., the number of overlapping accessions with phenotype and transcriptome data is different in different phenotypic datasets). In addition, we conducted TWAS with transcriptome data with (log) or without (non-transformation) a logarithm transformation. Finally, we re-distributed the raw TPM value for each gene into a consecutively numeric range from 0 to 2 as described in the GAPIT manual for TWAS with the implementation of MLM in GAPIT (Lipka *et al*., 2012), using a relatedness matrix based on gene expressions to correct the population structure.

### Plant materials and growth conditions

For the functional validation of the candidate genes, 34 T-DNA mutants covering 17 candidate genes associated with TWAS traits were purchased from the Arabidopsis Biological Resource Center (ABRC) and Nottingham Arabidopsis Stock Centre (Table S1). Homozygous lines were isolated using genotyping.

After three days of stratification in the dark at 4°C on agar plates containing half-strength Hoagland solution with 1% sucrose (Chiou *et al*., 2006), seeds were germinated and grown in controlled growth chambers at the following settings: 16 h light/8 h darkness, 16°C constant temperature, 60-80% humidity (Grimm *et al*., 2017) with white light at 120–150 μE m^−2^ sec^−1^. Four replicates of fourteen-day-old seedlings were transferred to pots and placed in a randomized block design. To minimize position effects, all pots within a tray were rotated 180°C and the tray order was shuffled from the first to the last every other day. Flowering time was scored as days until the inflorescence stem elongated to 1 cm.

### Identification of expression QTL

The expression QTL (eQTL) was identified using the same 631 accession inbred lines analyzed for TWAS of flowering at 16°C. To determine eQTLs involved in flowering regulation, individual raw TPMs of the selected trait-associated genes were taken as expression traits for the GWAS analysis. GWAS was conducted in GWA-Portal (https://gwas.gmi.oeaw.ac.at/) against an imputed full sequence dataset, which contained 2,029 accessions with missing data imputed (The 1001 Genomes Consortium, 2016), using MLM (named as an accelerated mixed model [AMM] in GWA-Portal) (Seren *et al*., 2012). MLM considers kinship to reduce the confounding effects of population structure (Kang *et al*., 2008; Seren *et al*., 2012; The 1001 Genomes Consortium, 2016). The significant SNPs (-log_10_[*P*] ≥ 6 and MAC ≥ 10) for each gene were grouped into clusters (1 Mb region) and shown by a heatmap.

## Results

### Potential of TWAS analysis to identify causative genes

We downloaded the transcriptome data of rosette leaves from 727 Arabidopsis natural accessions (Kawakatsu *et al*., 2016) and phenotype datasets from AraPheno (https://arapheno.1001genomes.org/). For the transcriptomes, after quality control, mapping, quantification, and elimination of chloroplast and mitochondria genes, 36,631 genes were retained. For TWAS, we removed genes with zero or scarce variance in expression level across more than 95% of the lines in the transcriptomic dataset using the nearZeroVar function from the R caret package. Thus, the number of explanatory variables associated with individual traits may differ. In addition, transcriptome data with or without logarithm (log) transformation were tested for the association. For the phenotype, we selected four comprising more than 100 accessions with both phenotypes and transcriptome data to ensure statistical power for the association. Finally, association mapping was performed using MLM in an R package Genome Association and Prediction Integrated Tool (GAPIT) (Lipka *et al*., 2012).

We first analyzed the growth allometry traits (study 31 in AraPheno), in which growth scaling- and fitness-related traits (growth rate, relative growth rate, fruit number, scaling exponent and rosette dry mass) of 451 sixteen-day-old Arabidopsis accessions were measured (Vasseur *et al*., 2018). Among them, 270 accessions having both traits and transcriptome data were applied for TWAS. Based on the analysis using non- or log-transformed expression values, several genes showing a significant correlation (false discovery rate [FDR] ≤ 0.05) with these traits were identified (Table S2). Among them, we noticed several known flowering regulators (Bouché *et al*., 2016; Brachi *et al*., 2010; Huo *et al*., 2016) associated with more than one trait (Fig. 1, Supplementary Table S2). For example, *SOC1* and *FLC* are associated with traits of “growth rate”, “fruit number”, “scaling exponent”, and “rosette dry mass”‘. *AGAMOUS-LIKE 24* (*AGL24*) is associated with traits of “growth rate” and “rosette dry mass”, while *AGL8* is associated with traits of “growth rate” and “scaling exponent”. This finding suggests that flowering time is a crucial parameter for allometric diversification, in agreement with the previous study showing that resource-acquisitive plants (i.e., early-flowering/fast-growing ecotypes) are more adapted to hotter and drier regions (Vasseur *et al*., 2014). It is worth mentioning that although a link between flowering time and growth allometry was proposed in the previous study, no association was identified (Vasseur *et al*., 2018). Our TWAS identified flowering genes for controlling growth allometry, providing a potential connection between these two traits.

**Fig. 1.**
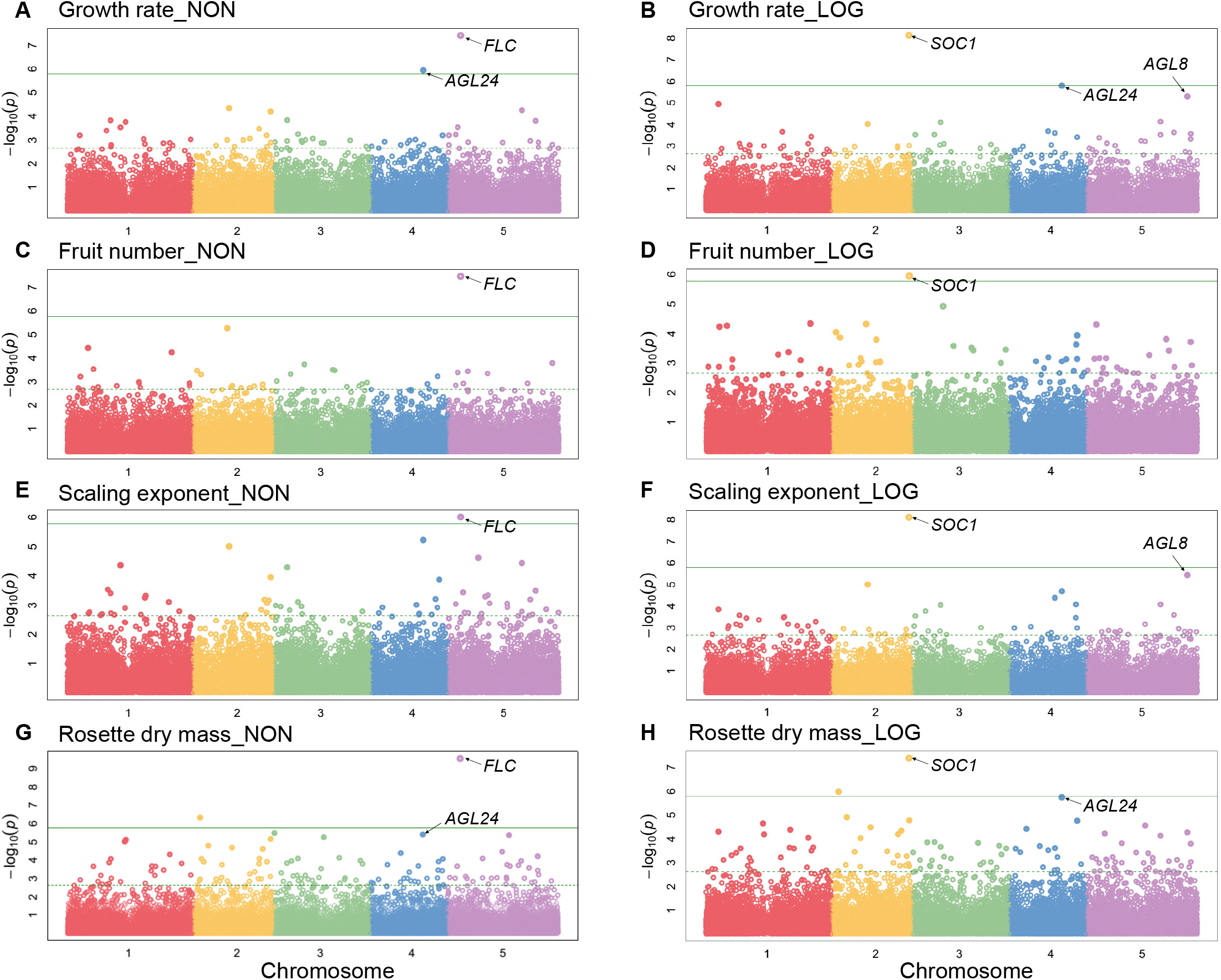
TWAS of growth allometry-related traits. Manhattan plots of TWAS results for growth allometry-related traits, in which non (NON)-transformed (A, C, E, G) and log (LOG)-transformed (B, D, F, H) gene expression values were used for the association. Only flowering-related genes showing significant (FDR ≤ 0.05) association with indicated phenotypes were labeled. The green solid and dashed lines represent the threshold of Bonferonni and FDR significance, respectively, provided by GAPIT.

Next, we analyzed the leaf metabolites (study 4 in AraPheno) of four-week-old Arabidopsis generated from liquid chromatography-mass spectrometry (LC-MS) (Strauch *et al*., 2015). Among 440 accessions, 127 accessions having both traits and transcriptome data were applied for TWAS. Remarkably, *D-AMINO ACID RACEMASE1* (*DAAR1*) was identified as the top candidate gene associated with the abundance of N-Malonyl-D-allo-Isoleucine represented at M130T666, M172T666, and M216T665 (M = mass-to-charge ratio; T = retention time in LC-MS) regardless of whether non-transformed or log-transformed data were analyzed (Fig. 2). This gene has also been identified in the previous GWAS, followed by a functional validation using a T-DNA insertional mutant (Strauch *et al*., 2015), thereby supporting the promise of our TWAS analysis to determine the causative genes.

**Fig. 2.**
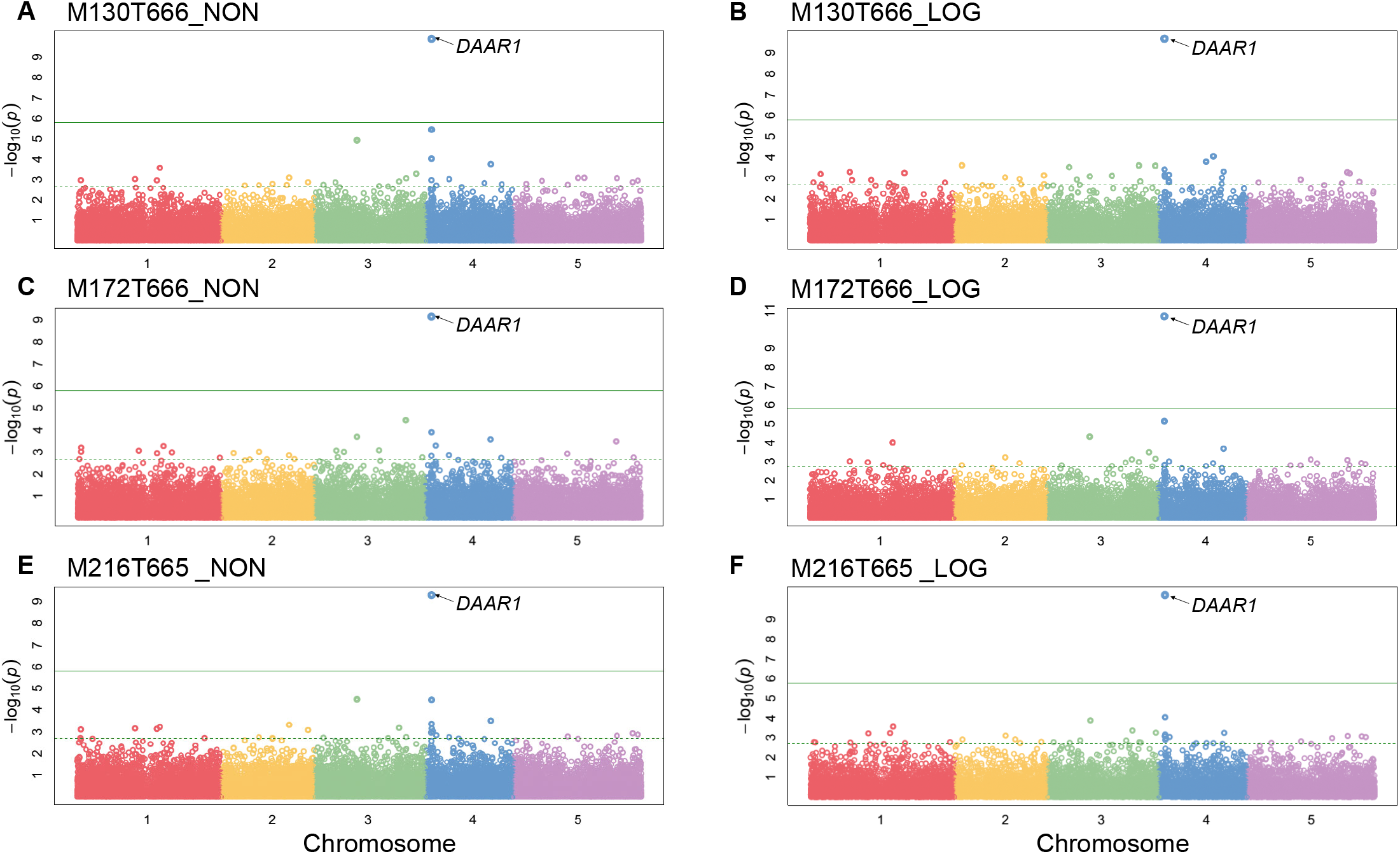
TWAS of the metabolite trait. Manhattan plots of TWAS results for the production of a metabolite N-Malonyl-D-allo-Isoleucine, identified at M130T666 (A and B), M172T666(C and D), and M216T665 (E and F) (M = mass-to-charge ratio; T = retention time in LC-MS), while (A), (C) and (E) using non (NON)-transformed, and (B), (D) and (F) using log (LOG)-transformed gene expression values for the association. The green solid and dashed lines represent the threshold of Bonferonni and FDR significance, respectively, provided by GAPIT.

### TWAS analysis of flowering time

In AraPheno (study 12), a dataset of flowering phenotype, “days to flowering” (The 1001 Genomes Consortium, 2016) was used for the analysis. Among more than 1,100 accessions, 649 and 631 having both traits and transcriptome data of flowering at 10°C and 16°C, respectively, were applied for TWAS. As shown in Figure 3A and 3B, the flowering time at 10°C and 16°C is distributed normally (Shapiro–Wilk score [-log_10_(*P*)]: 10.17 for 10°C and [-log_10_(*P*)]: 11.98 for 16°C), suggesting that, as reported previously, flowering time is polygenetically regulated (The 1001 Genomes Consortium, 2016).

**Fig. 3.**
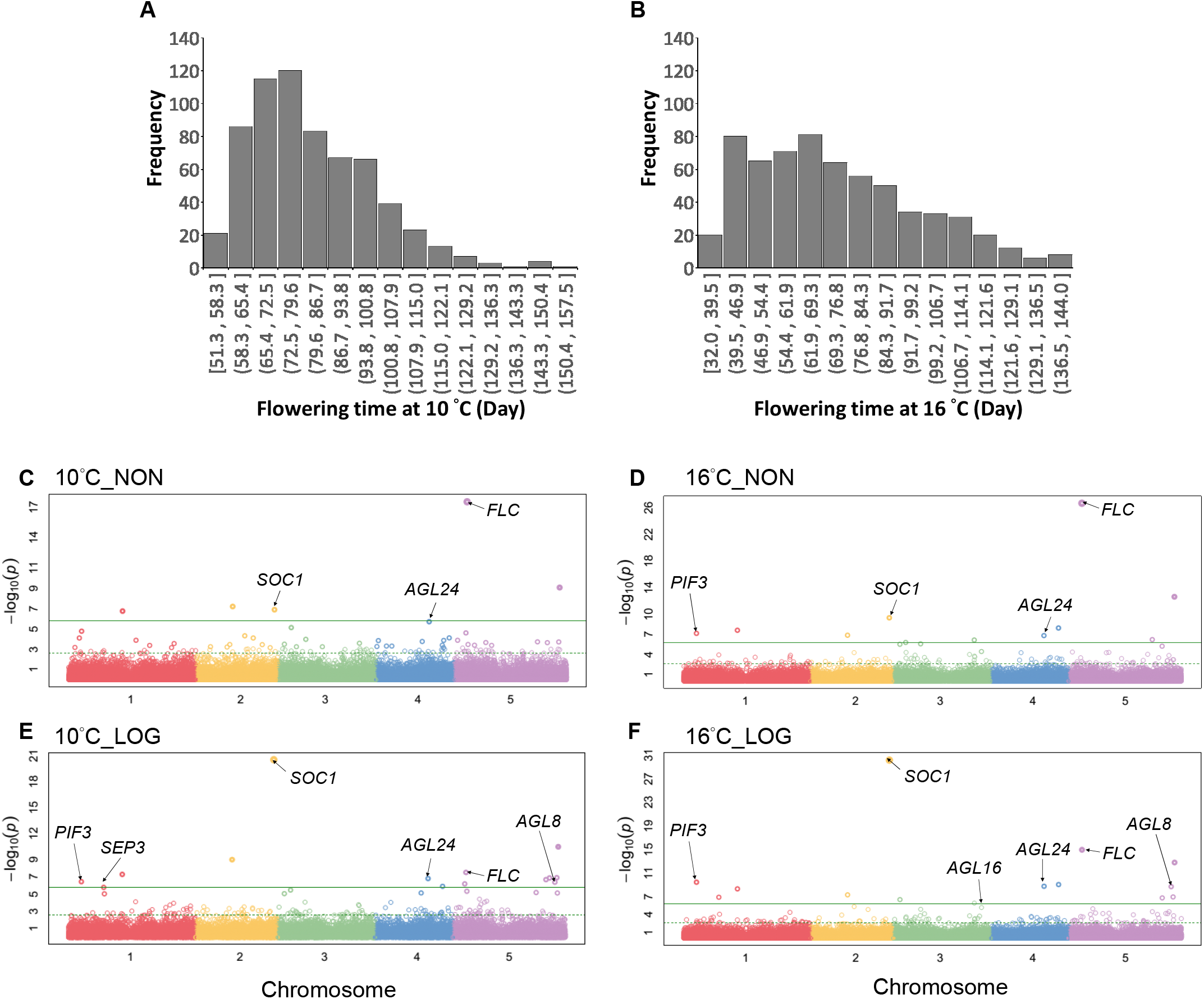
Associations for flowering time trait. Phenotypic distribution of flowering time at 10°C (A) and 16°C (B). Manhattan plots of TWAS results for the flowering time at 10°C (C and E) and 16°C (D and F) while (C) and (D) using non (NON)-transformed, and (E) and (F) using log (LOG)-transformed gene expression values for the association. TWAS was analyzed using GAPIT, in which green solid and dashed lines indicate the thresholds of Bonferonni and FDR significance, respectively.

In the TWAS result (Fig. 3C-F, Supplementary Table S3), many *a priori* flowering regulators (Bouché *et al*., 2016; Brachi *et al*., 2010) associated with the flowering time at 10°C or 16°C were evident because of their high significant (FDR ≤ 0.05) correlations. *PHYTOCHROME INTERACTING FACTOR 3* (*PIF3*), *SOC1, AGL24*, and *FLC* were detected using non-transformed expression values (Fig. 3C, D, Supplementary Table S3), and *SEPALLATA3* (*SEP3*), *AGL8* and *AGL16* were additionally detected using log-transformed expression values (Fig. 3E, F, Supplementary Table S3). Moreover, we found that the expression level of *FLC* showed a significantly (*P* ≤ 0.05) high degree of correlation (0.5 ≤ |Pearson’s r| ≤ 1) with the flowering time at 10 and 16°C. Expression levels of *SOC1, PIF3, SEP3*, and *AGL8* showed a moderate degree of correlation (0.3 ≤ |Pearson’s r| ≤ 0.49), and expression levels of *AGL16* and *AGL24* showed a low degree of correlation (0 < |Pearson’s r| ≤ 0.29) to flowering time.

### Functional validation of TWAS-identified flowering candidate genes

With a significant cutoff of FDR at 0.05, we identified 24 genes associated with flowering at 10°C (17 genes) or 16°C (21 genes) and 14 genes for both (combined genes listed in results of “non-transformation” and “log-transformation”, Table 1) by TWAS. Furthermore, when we analyzed another flowering-time-related dataset (Grimm *et al*., 2017), except for *AT5G19140* (*AILP1*), the other 23 genes were also identified (Supplementary Table S4). This result reinforces the potential role of these genes in flowering-time control.

**Table 1:** Twenty-four Arabidopsis genes associated with flowering via TWAS.

| AGI* | Gene name | FT 16°C<sup>a</sup> |  | FT 10°C<sup>a</sup> |  |
| --- | --- | --- | --- | --- | --- |
|  |  | Non-transformation | Log-transformation | Non-transformation | Log-transformation |
| AT5G10140 | FLC | 6.30E-23 | 1.54E-11 | 9.07E-14 | 2.31E-04 |
| AT2G45660 | SOC1/AGL20 | 3.11E-06 | 1.99E-26 | 9.42E-04 | 8.56E-17 |
| AT1G09530 | PIF3/PAP3 | 3.34E-04 | 2.37E-06 | ns | 1.08E-03 |
| AT4G24540 | AGL24 | 5.45E-04 | 7.64E-06 | 9.51E-03 | 5.88E-04 |
| AT5G60910 | AGL8 | ns | 7.64E-06 | ns | 1.12E-03 |
| AT3G57230 | AGL16 | ns | 1.17E-02 | ns | ns |
| **AT5G63120** | RH30 | 3.31E-09 | 1.45E-09 | 1.24E-05 | 4.97E-07 |
| **AT1G35180** | TRAM, LAG1 and CLN8 (TLC)<br>lipid-sensing domain<br>containing protein | 1.38E-04 | 1.59E-05 | 1.04E-03 | 3.10E-04 |
| **AT2G20440** | Ypt/Rab-GAP domain of gyp1p<br>superfamily protein | 5.45E-04 | 1.47E-04 | 6.11E-04 | 1.01E-05 |
| AT5G55120 | VTC5 | 1.21E-02 | 3.59E-04 | ns | 6.93E-04 |
| **AT5G62165** | AGL42/FYF | ns | 2.98E-04 | ns | 5.74E-04 |
| AT4G33625 | vacuole protein | 7.87E-05 | 4.78E-06 | ns | 2.87E-03 |
| AT5G49360 | BXL1 | 1.93E-03 | 4.44E-02 | ns | 1.14E-02 |
| AT3G09920 | PIP5K9 | 4.23E-03 | ns | ns | ns |
| AT3G52480 | transmembrane protein | 2.00E-03 | 2.38E-03 | ns | ns |
| **AT3G05670** | RING/U-box protein | 6.76E-03 | 6.52E-04 | ns | 1.30E-02 |
| AT3G19150 | ACK1, KRP6 | 6.26E-03 | 2.10E-02 | ns | ns |
| AT1G23870 | TPS9 | ns | 3.01E-04 | ns | 3.52E-03 |
| AT5G59590 | UGT76E2 | ns | 4.44E-02 | ns | ns |
| AT5G17460 | DFR1 | ns | 1.75E-02 | ns | ns |
| AT2G32290 | BAM6/BMY5 | ns | 2.11E-02 | ns | ns |
| **AT5G19140** | AILP1 | ns | ns | ns | 5.08E-02 |
| AT5G09300 | thiamin diphosphate-binding<br>fold (THDP-binding)<br>superfamily protein | ns | ns | ns | 1.61E-03 |
| AT5G57340 | ras guanine nucleotide<br>exchange factor Q-like protein | ns | ns | ns | 5.74E-04 |

<sup>a</sup>indicates phenotypes studied in The 1001 Genomes Consortium, 2016
The values indicate FDR-adjusted P values provided by GAPIT according to either log-transformed or non-transformed transcriptome data.
FT: flowering time
ns, not significant (FDR > 0.05).
\*The genes inside the gray area are the six known flowering genes, and the genes marked in bold are six previously unknown flowering-related genes identified in this work (see Fig. 4).

Among the 24 TWAS-identified candidate genes, six (*FLC, SOC1, PIF3*, and *AGL8*/*16*/*24*) are well-characterized flowering genes (Bouché *et al*., 2016; Brachi *et al*., 2010). We next focused on functional validation of the rest uncharacterized candidate genes by analyzing 27 available T-DNA insertional mutants (Supplementary Table S1). Because most candidate genes were identified from 16°C, we thus assessed flowering time at 16°C using the number of days until the first inflorescence stem elongated to one centimeter as the phenotype equivalent to the “days to flowering (DTF) 2” in Grimm et al. (2017). We considered genes as potential regulators when the expression level and flowering time in their corresponding mutants showed significant differences compared to the wild-type (WT) Columbia-0 (Col-0) control. Using this criterion, we found that loss-of-function in three genes, *AT1G35180* (LAG1 and CLN8 [TLC] lipid-sensing domain-containing protein; TRAM), *AT3G05670* (RING/U-box protein), and *AT5G63120* (*RNA HELICASE 30, RH30*), resulted in a significant delay in flowering time (1.4–5.2 days) (Fig. 4). On the other hand, loss-of-function in another two genes, *AT2G20440* (Ypt/Rab-GAP domain of gyp1p superfamily protein) and *AT5G19140* (*AILP1*), promoted flowering by 1.3–1.9 days (Fig. 4). Intriguingly, the insertion of T-DNA in the 5’ untranslated region of *AT5G62165* (*AGL42*/*FOREVER YOUNG FLOWER, FYF*) resulted in increased expression accompanied by promoting flowering for 2.55 days (Fig. 4). Out of 24 candidate genes from our TWAS, we confirmed six with previously uncharacterized functions to flowering time in addition to the six known flowering genes.

**Fig. 4.**
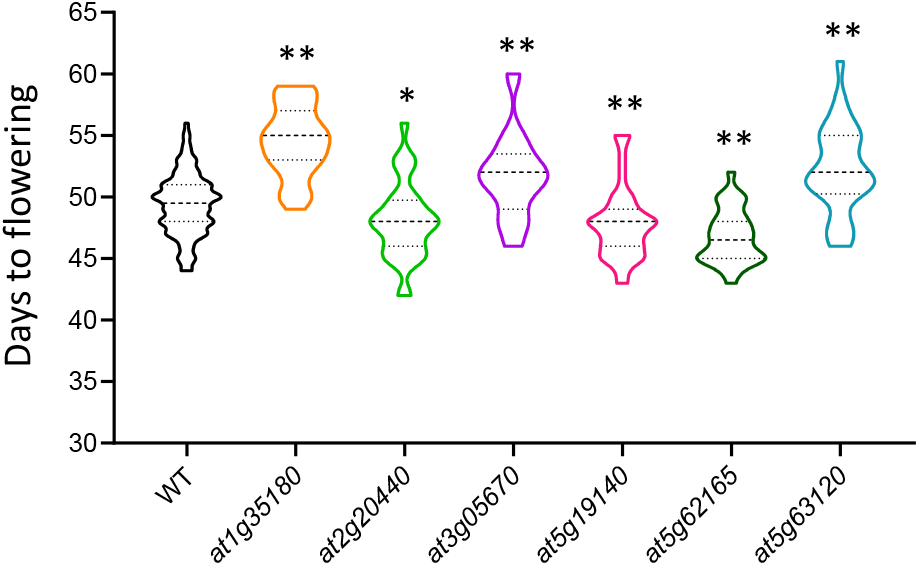
Functional validation of TWAS identified candidate genes. T-DNA insertional mutants of the following genes were examined: TRAM, LAG1 and CLN8 (TLC) lipid-sensing domain-containing protein (*AT1G35180*; SALK_128954C), Ypt/Rab-GAP domain of gyp1p superfamily protein (*AT2G20440*; SALK_006098C), RING/U-box protein (*AT3G05670*; WiscDsLoxHs229_04H); *AILP1* (*AT5G19140*; WiscDsLox393-396N12), *AGL42/FYF* (*AT5G62165*; SALK_076684C), and *RH30* (*AT5G63120*; WiscDsLoxHs044_01H). n = 32 biologically independent samples for the mutants, while in wild-type (WT) Col-0, n = 220. The gray dotted lines within the violin plots represent the 25th and 75th percentiles, while the black dashed lines represent the median. The *P* value was determined by two-sided Student’s *t*-test (*, *P* < 0.05; **, *P* < 0.01).

### eQTL analysis integrates genetic factors into the TWAS result

To further explore upstream genetic factors governing the transcript abundance of trait-associated genes among accessions, we performed eQTL analysis implemented in GWA-Portal (Seren, 2018; https://gwas.gmi.oeaw.ac.at/) of 24 TWAS-identified flowering candidate genes against an imputed full sequence dataset. We identified 496 SNPs passing the threshold: -log_10_(*P*) ≥ 6, minor allele frequency (MAF) ≥ 0.05, and minor allele count (MAC) ≥ 10 as significant eQTLs (Fig. 5A, Supplementary Table S5). We detected *cis*-eQTLs in six TWAS-identified candidate genes, *PIF3, PHOSPHATIDYL INOSITOL MONOPHOSPHATE 5 KINASE* (*PIP5K9*), *ARABIDOPSIS CDK INHIBITOR 1*/*KIP-RELATED PROTEIN 6 (*ACK1/*KRP6*/*ICK4*), a transmembrane protein *AT3G52480, FLC*, and *AILP1* (indicated by black arrows in Fig. 5A). Within the 1 kb region, 82, 7, 8, 6, 22, and 203 significant SNPs were associated with the corresponding genes, respectively. Moreover, one cluster of six significant SNPs (262,690, 298,815, 299,748, 304,857, 305,713, and 317,979 bp) on Chr 4 was detected as a *trans*-eQTL (labeled on top of Fig. 5A by a red star) for four flowering regulators: *FLC, AGL8, SOC1*, and a TWAS newly identified *AGL42* (Fig. 5B-E). Notably, this *trans*-regulatory hotspot covered 14 genes, including the whole gene bodies of two known flowering-related genes, *FRI* (Chr 4: 269,026 - 271,503, from start to stop codon) and *UB-LIKE PROTEASE 1B* (*ULP1B*; *AT4G00690*; Chr 4: 281,313 – 283,129, from start to stop codon; Brachi *et al*., 2010). Thus, a combination of TWAS and eQTL analyses further revealed the genetic connection linking transcripts of flowering regulators to the phenotypic outcome.

**Fig. 5.**
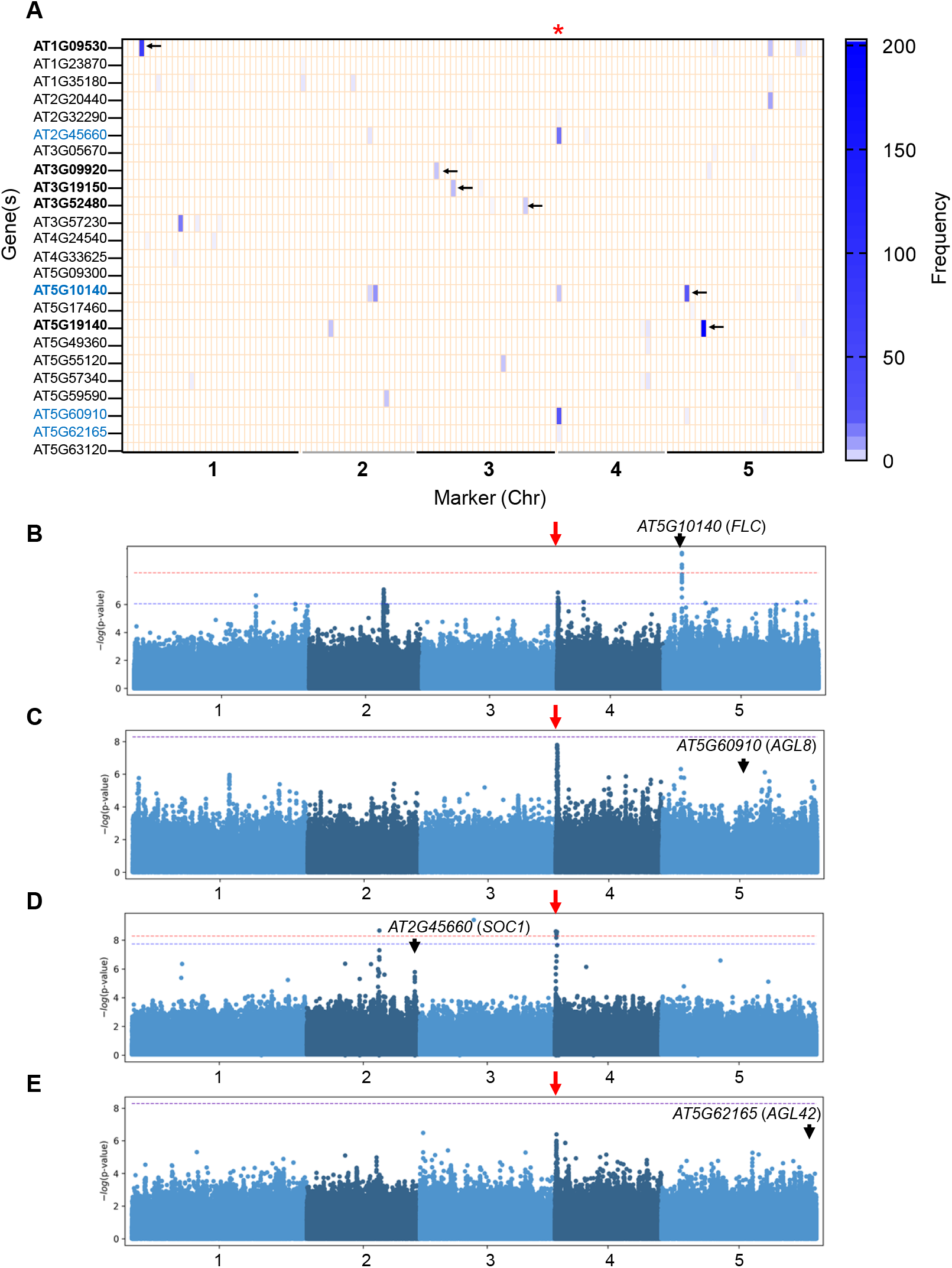
eQTL analysis integrates the information of genetic factors into the TWAS result. (A) A heatmap shows the physical position of TWAS-identified candidate genes on the vertical axis against their significantly associated SNPs on the horizontal axis. Each cell represents a genomic region of 1 Mb wherein a gradient color indicates the number of significant SNPs (namely frequency). *Cis*-eQTLs (indicated by black arrows) are identified for six genes, *AT1G09530* (*PAP3*/*PIF3*), *AT3G09920* (*PIP5K9*), *AT3G19150* (*ACK1*/*KRP6*/*ICK4*), a transmembrane protein *AT3G52480, AT5G10140* (*FLC*), and *AT5G19140* (*AILP1*), marked in bold. A *trans*-eQTL hotspot located on the top of Chr 4 (indicated by a red star) is identified to be associated with four genes, *AT2G45660* (*SOC1*), *AT5G60910* (*AGL8*), *AT5G10140* (*FLC*), and *AT5G62165* (*AGL42/FYF*), marked in blue. (B-E), Manhattan plots present GWAS results for the expression levels of four TWAS-identified candidate genes mentioned above. The black arrow marks their physical locations on Manhattan plots. The significant peak on Manhattan plots representing the *trans*-eQTL hotspot in (A) was indicated by a red arrow. These analyses were conducted using GWA-Portal. Thresholds for Bonferroni (red dashed line) and Benjamini Hochberg (blue dashed line) correction provided by GWA-Portal are indicated.

### Haplotypes on Chr4 discriminate flowering-related responses across different subgroups of accessions

To clarify the connection between the trans-regulatory hotspot and flowering time, we first arranged allelic combinations of the six significant SNPs. Due to high linkage disequilibrium, only eight haplotypes, ‘TCAGTA’, ‘TCAGTT’, ‘TCAATA’, ‘TCAACA’, ‘CTGACT’, ‘CTGATA’, ‘CCAGTA’, and ‘TTAGTA’, denoted as Hap1 to Hap8, respectively, were identified (Fig. 6A; left panel). Among these eight haplotypes, only the Hap1 and Hap5 groups had sufficient sample sizes (more than ten accessions) and were selected for functional characterization. As mentioned, these haplotypes covered gene bodies of two flowering-related genes, *FRI* and *ULP1B*. Because of the crucial role of *FRI* in regulating downstream gene expression, especially *FLC*, we then focused on *FRI* in the subsequent analysis.

**Fig. 6.**
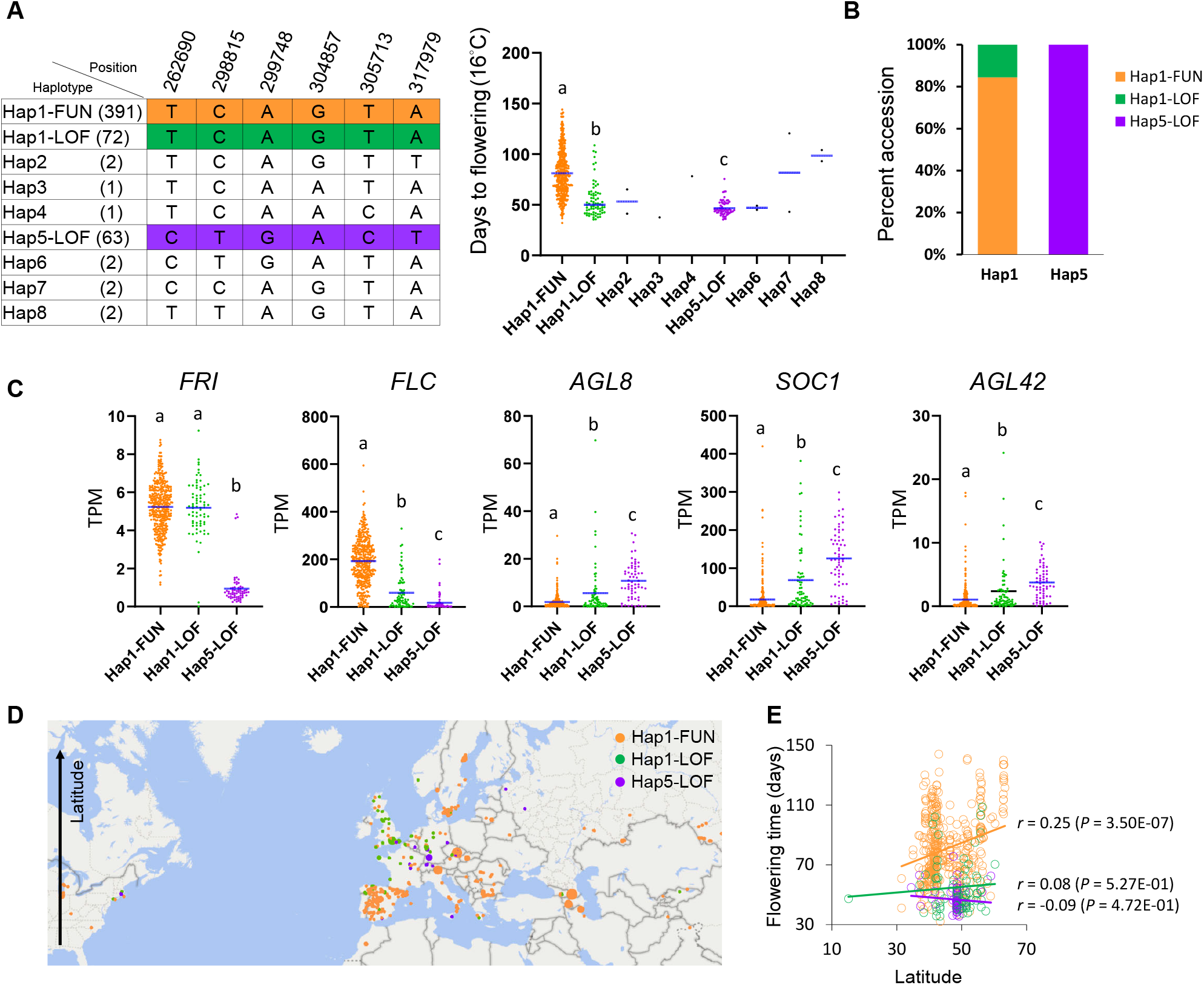
Haplotype covering the *FRI* gene body connects expressions of several flowering regulators with flowering time in Arabidopsis accessions. (A) The six significantly associated SNPs composed of the *trans*-eQTL hotspot on Chr 4 result in eight haplotypes. Values in the brackets indicate the number of accessions in each subgroup (left panel). In the dot plot (right panel), the blue line inside a cluster of dots indicates the mean of phenotypic values. FUN = functional, and LOF = Loss-of-functional, *FRI* alleles. (B) Percentage of FUN- or LOF-*FRI* alleles in Hap1 and Hap5 haplogroups. (C) Expression levels of *FRI* and another four TWAS-identified flowering genes, including *FLC, AGL8, SOC1*, and *AGL42*, in different haplogroups. Different letters above the bars indicate statistical differences from one-way ANOVA followed by the least significant difference test (LSD) at a probability of *P* < 0.05. (D) Global distribution of different genotypic accessions. Symbols in different colors are the haplogroups as indicated. (E) Correlation between flowering time (16°C) and latitude among different subgroups of accessions.

Zhang and Jiménez-Gómez, (2020) recently defined 103 distinct *FRI* alleles based on nonsynonymous changes in 1,016 Arabidopsis accessions. Most accessions that do not require vernalization to induce flowering carry non-functional alleles of *FRI*. Two of the most common cases are the laboratory strains Col-0 and Landsberg *erecta* (L*er*). *FRI*-Col-0 allele contains a 16-bp deletion, resulting in a premature stop codon (Johanson *et al*., 2000). *FRI*-L*er* allele carries a 376-bp deletion combined with a 31-bp insertion in the promoter region that removes its translational start and dramatically reduces its expression (Schmalenbach *et al*., 2014). Accordingly, we integrated the information about the *FRI* function (Zhang and Jiménez-Gómez, 2020) into our analysis (Fig. 6B). In the Hap1-accessions, 84.4% have functional *FRI* (Hap1-FUN), but 15.6% have loss-of-function *FRI* (Hap1-LOF), in which 26.4% (19 out of 72) Hap1-LOF accessions carry the *FRI*-Col-0 allele. In Hap5, all accessions carry the loss-of-function *FRI*-L*er* allele, denoted as Hap5-LOF (Fig. 6B). The late-flowering accessions typically possess functional *FRI* and *FLC* genes (Johanson *et al*., 2000; Michaels and Amasino, 1999). As expected, Hap1-FUN accessions flowered the latest (mean of the number of days to flowering = 81), but Hap5-LOF accessions flowered the earliest (mean of the number of days to flowering = 47) among all accessions (Fig. 6A, right panel). The flowering time of Hap1-LOF accessions (mean of the number of days to flowering = 55) was close to Hap5-LOF (Fig. 6A, right panel).

We also found that the *FRI* expression is distinct in Hap1 and Hap5. The *FRI* expression level is higher in Hap1 accessions than in Hap5 (Fig. 6C). However, *FRI* expression levels in Hap1-FUN and Hap1-LOF showed no significant difference. This is probably because nonsynonymous changes in the *FRI* allele of most Hap1-LOF do not affect its transcriptional abundance. Similarly, Hap1-FUN accessions had the highest expression level of *FLC*, while Hap5-LOF accessions had the lowest (Fig. 6C). The expression of *FLC* changed along with the expression of functional *FRI* (Hap1-FUN vs. Hap5-LOF) further strengthens the previous finding that *FLC* acts synergistically with *FRI* to repress flowering (Johanson *et al*., 2000; Koornneef *et al*., 1994; Sheldon *et al*., 2000). On the contrary, Hap1-FUN accessions have significantly lower expressions of *AGL8, SOC1*, and *AGL42* than Hap1-LOF and Hap5-LOF accessions (Fig. 6C). This observation agrees with the report on the suppression of *SOC1* by *FLC* (Michaels and Amasino, 2001). Overall, our findings provide additional evidence supporting the importance of the *FRI-FLC-SOC1* signaling module in regulating flowering time from the perspective of genetic variations.

From the geographic distribution, accessions carrying Hap1-LOF and Hap5-LOF seem to grow mainly in Northwest Europe (Fig. 6D). In line with the previous report showing a strong correlation between flowering time and latitude among the accessions with functional alleles of *FRI* (Stinchcombe *et al*., 2004), we also found that the flowering time of Hap1-FUN accessions was correlated with latitude (Fig. 6E; Pearson’s r = 0.25, *P* = 3.50E-07). However, this correlation disappeared in the accessions carrying Hap5-LOF (Pearson’s r = -0.09, *P* = 4.72E-01) and Hap1-LOF (Pearson’s r = 0.08, *P* = 5.27E-01) (Fig. 6E). This result indicates that an association between flowering time and latitude is only detectable under a functional FRI background.

In summary, our analyses reveal the haplotype on Chr 4 bridges the function of *FRI* in controlling downstream flowering regulators and the variation in flowering time, resulting in plant adaptation to different habitats.

## Discussion

### Feasibility of TWAS

This study employed TWAS by combining transcriptomes and phenotypes from Arabidopsis 1,001 Genomes to analyze gene-trait associations. We identified several trait-associated genes for various phenotypes. For example, we revealed that flowering regulatory genes are predictive of allometric diversification (Fig. 1, Supplementary Table S2) and the *DAAR1* gene produces the metabolite of N-Malonyl-D-allo-Isoleucine (Fig. 2); we also revealed some novel flowering regulators (Fig. 3, 4). TWAS thus appears to be a powerful tool for identifying candidate genes even though the transcriptome data is generated in a material different from phenotyping, demonstrating the feasibility and the potential value of TWAS. Additionally, compared with GWAS, which alone often poses challenges in pinpointing causative genes within an associated LD-decay region, TWAS was more straightforward in prioritizing candidate genes.

### Discovery of additional flowering regulators

While we were conducting this TWAS analysis, two studies also reported the application of TWAS in Arabidopsis using a linear regression model (Lan *et al*., 2021) or an MLM for association mapping (Li *et al*., 2021). Since Li et al. (2021) and our group both conducted TWAS for the flowering time at 16°C (631 samples) using GAPIT, we naturally identified some genes in common (FDR ≤ 0.05), including four flowering regulators, *SOC1, FLC, PIF3*, and *AGL24*, and five potential candidate genes: *ACK1, RH30, AT2G20440, AT1G35180*, and *AT5G49360*. However, there were some differences: we identified *AGL8* and *AGL16*, while Li et al. (2021) identified *GA INSENSITIVE DWARF1C* (*GID1C*). In addition, we identified an additional 13 candidate genes (Table 1). We reason that the difference may be due to the parameter setting for transcriptome data processing; furthermore, whether expression levels are transformed may matter. For the potential candidates, Lan et al. (2021) also found that *AT4G20440* and *RH30* were negatively correlated to flowering time. However, we noticed that the threshold for gene-trait connection in simple correlation applied in the study of Lan et al. (2021) is not stringent (|Pearson’s r| > 0.3 and |Spearman’s r| > 0.3). Even so, fewer flowering genes (7 *a priori* flowering-related genes found at 10°C and 15 genes at 16°C based on the same dataset from Arapheno) were identified when a simple correlation rather than a model of MLM was used, suggesting that the result can be improved if residuals, such as population structure and kinship, are taken into account.

Furthermore, we provided functional validation for candidate genes (Fig. 4). In addition to six known flowering genes, we identified six previously uncharacterized flowering regulators. They are *AT1G35180* encoding a TRAM, LAG1 and CLN8 (TLC) lipid-sensing domain-containing protein, *AT2G20440* encoding a Ypt/Rab-GAP domain of gyp1p superfamily protein, *AT3G05670* encoding a RING/U-box protein, *AILP1* encoding an aluminum-induced protein with YGL and LRDR motifs, *AGL42/FYF* encoding a MADS box transcription factor, and *RH30* encoding a P-loop containing nucleoside triphosphate hydrolases superfamily protein. These genes are novel flowering-related genes. There are no prior studies directly linking these genes to flowering time except *FYF* and RNA helicase genes. FYF is known as a repressor controlling floral organ senescence and abscission (Chen *et al*., 2011), and mutations on the DEAD-box of RNA helicase genes influence flowering time in Arabidopsis probably due to the defect of RNA processing (Bush *et al*., 2015; Gong *et al*., 2005; Herr *et al*., 2006; Mahrez *et al*., 2016). Moreover, based on the prediction by ATTED II (Obayashi *et al*., 2018), *RH30* is highly co-expressed with *AT2G20440*, another flowering-related gene identified in our TWAS. The functional validation of associated candidate genes offers an evaluation of the feasibility of our TWAS method.

It is worth noting that six flowering-related genes, including three well-characterized (*FLC, SOC1* and *AGL24*) and three previously unknown (*AT1G35180, AT2G20440* and *RH30*) genes, were identified in all approach combinations (i.e., with non- and log-transformed values at 10°C or 16°C, Table 1). They were functionally validated (Fig. 4) and highly ranked among the 24 candidate genes (Table 1).

### Haplotype that covered the FRI gene body links flowering-regulatory genes to flowering time

Our analysis found two haplotypes on the top of Chr4, Hap1 and Hap5, which discriminated flowering phenotype significantly (Fig. 6). Most Hap1-accession have functional FRI (Hap1-FUN), but the rest carry loss-of-function FRI (Hap1-LOF), one-fourth of which have the *FRI*-Col-0 allele. In Hap5-LOF, all accessions carry the *FRI*-L*er* allele (Fig. 6B). Because of the null function of FRI, Col-0 and L*er* exhibit early flowering (Johanson *et al*., 2000; Schmalenbach *et al*., 2014). Indeed, Hap1-LOF (some are *FRI*-Col-0) and Hap5-LOF accessions (*FRI*-L*er*) flower significantly earlier than Hap1-FUN accessions (Fig. 6A). Intriguingly, this observation implies that the emergence of the Hap1-LOF subgroup from Hap1 may occur after the Hap1-Hap5 split event. Our further analysis of gene expression reinforces the scenario that FLC acts synergistically with FRI to repress flowering in late-flowering accessions (Johanson *et al*., 2000; Koornneef *et al*., 1994; Sheldon *et al*., 2000) and suppression of *SOC1* expression by *FLC* (Fig. 6C) (Michaels and Amasino, 2001). Finally, the analysis of Hap1 and Hap5 haplotypes provided a genetic connection linking flowering gene expressions to flowering time. Multiple *FLC* haplotypes were previously found to underpin life history diversity in Arabidopsis (Li *et al*., 2014). However, given that the effect of variation in *FLC* can only be observed in the presence of a functional allele of *FRI* (Caicedo *et al*., 2004), we exposed the connection of a known flowering regulator FRI to the control of flowering time across different haplogroups of natural accessions. Of note, with the functional annotation by *FRI* allele, we revealed a recent split event during evolution; even though they are in the same haplogroup (e.g., Hap1), subgroups of natural accessions (Hap1-LOF vs. Hap-FUN) flower distinctly. The principle of our approaches, taking flowering traits as an example, is illustrated in Figure 7, explicating the connections across different datasets and the analysis outcomes.

**Fig. 7.**
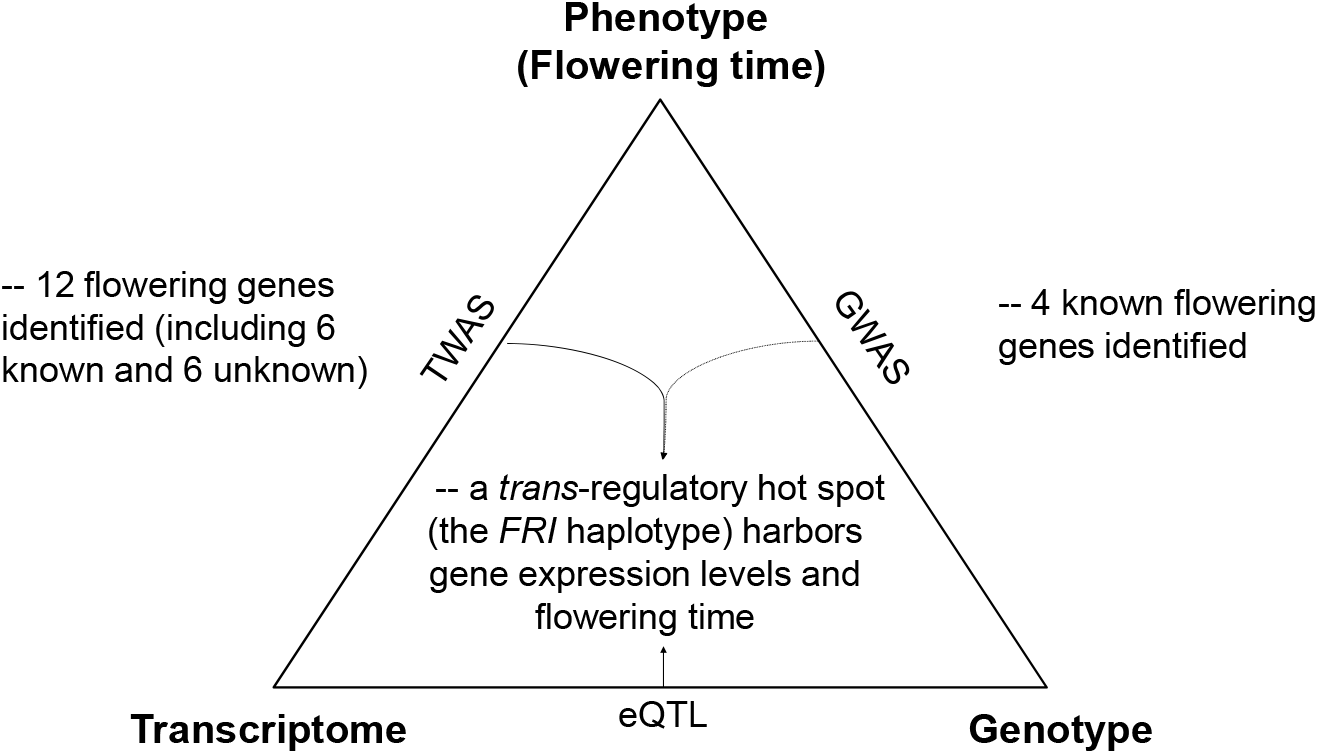
Illustration of the integrative approach used to identify genes related to flowering time. Information about Arabidopsis transcriptomes, genomes, and phenotypes from 1,001 Genomes allowed us to conduct GWAS, TWAS, and eQTL to reveal the gene/SNP-trait associations for flowering time. Integration of eQTL for TWAS identified candidate genes further revealing the *FRI* haplotype as a genetic harbor linking flowering genes to flowering time.

Our processing procedure for TWAS offers a framework to systematically assess the effectiveness of the TWAS approach for studying quantitative traits. In particular, for those plant species, such as cotton, rapeseed and garlic, with complex polyploid or huge genomes and for which a reference genome sequence is lacking (Chen *et al*., 2018; Harper *et al*., 2012; Havlickova *et al*., 2018; Li *et al*., 2020; Ma *et al*., 2021; Tang *et al*., 2020), TWAS provides a chance to detect potential trait-associated genes. In addition, our findings extend current knowledge by uncovering additional genetic regulators of flowering time and providing further evidence of the importance of the *FRI-FLC-SOC1* signaling module in regulating flowering time from the perspective of genetic variations.

## Supplementary data

**Supplementary Table S1**. List of T-DNA insertional mutants and the sequences of primer pairs for genotyping.

**Supplementary Table S2**. TWAS identified candidate genes associated with growth allometry-related traits (FDR ≤ 0.05) with (a) non- and (b) log-transformed data.

**Supplementary Table S3**. List of *a priori* flowering-related genes (FDR ≤ 0.05) under different TWAS conditions.

**Supplementary Table S4**. TWAS identified genes associated with flowering time.

**Supplementary Table S5**. Significant SNPs associated with TWAS-identified flowering genes (by GWA-Portal using MLM).

## Acknowledgements

We thank Drs. Hsin-Chou Yang and Yu-Jen Liang at the Institute of Statistical Science of Academia Sinica for constructive discussion to initiate this project. We also thank Ms. Su-Fen Chiang for genotyping T-DNA insertional mutants and Mr. Bo-Han Hou for troubleshooting programming editing.

## Author contributions

PSC, CRL, and TJC conceived and designed the study. PSC, PHC, and CRL performed computational analyses. PSC analyzed the data and performed the experiments. PSC, CRL, and TJC wrote and finalized the manuscript.

## Data availability

RNA-seq-derived transcriptomic data were downloaded from http://neomorph.salk.edu/1001.php (GSE80744). Phenotypic datasets and whole genome sequences of Arabidopsis were available in the AraPheno (https://arapheno.1001genomes.org/) and 1,001 Genomes (https://1001genomes.org/data/GMI-MPI/releases/v3.1/1001genomes_snp-short-indel_only_ACGTN.vcf.gz), respectively. The data that supports the findings of this study are available in the supplementary material of this article.

## References

Atwell S, Huang YS, Vilhjalmsson BJ, Willems G, Horton M, Li Y, Meng D, Platt A, Tarone AM, Hu TT, Jiang R, Muliyati NW, Zhang X, Amer MA, Baxter I, Brachi B, Chory J, Dean C, Debieu M, de Meaux J, Ecker JR, Faure N, Kniskern JM, Jones JD, Michael T, Nemri A, Roux F, Salt DE, Tang C, Todesco M, Traw MB, Weigel D, Marjoram P, Borevitz JO, Bergelson J, Nordborg M. 2010. Genome-wide association study of 107 phenotypes in Arabidopsis thaliana inbred lines. Nature 465, 627–631.

Blumel M, Dally N, Jung C. 2015. Flowering time regulation in crops-what did we learn from Arabidopsis? Curr Opin Biotechnol 32, 121–129.

Bolger AM, Lohse M, Usadel B. 2014. Trimmomatic: a flexible trimmer for Illumina sequence data. Bioinformatics 30, 2114–2120.

Bouché F, D’Aloia M, Tocquin P, Lobet G, Detry N, Perilleux C. 2016. Integrating roots into a whole plant network of flowering time genes in Arabidopsis thaliana. Sci Rep 6, 29042.

Brachi B, Faure N, Horton M, Flahauw E, Vazquez A, Nordborg M, Bergelson J, Cuguen J, Roux F. 2010. Linkage and association mapping of Arabidopsis thaliana flowering time in nature. PLoS Genet 6, e1000940.

Bush MS, Crowe N, Zheng T, Doonan JH. 2015. The RNA helicase, eIF4A-1, is required for ovule development and cell size homeostasis in Arabidopsis. Plant J 84, 989–1004.

Caicedo AL, Stinchcombe JR, Olsen KM, Schmitt J, Purugganan MD. 2004. Epistatic interaction between Arabidopsis FRI and FLC flowering time genes generates a latitudinal cline in a life history trait. Proc Natl Acad Sci U S A 101, 15670–15675.

Chen MK, Hsu WH, Lee PF, Thiruvengadam M, Chen HI, Yang CH. 2011. The MADS box gene, FOREVER YOUNG FLOWER, acts as a repressor controlling floral organ senescence and abscission in Arabidopsis. Plant J 68, 168–185.

Chen X, Liu X, Zhu S, Tang S, Mei S, Chen J, Li S, Liu M, Gu Y, Dai Q, Liu T. 2018. Transcriptome-referenced association study of clove shape traits in garlic. DNA Res 25, 587–596.

Cheng CY, Krishnakumar V, Chan AP, Thibaud-Nissen F, Schobel S, Town CD. 2017. Araport11: a complete reannotation of the Arabidopsis thaliana reference genome. Plant J 89, 789–804.

Chiou TJ, Aung K, Lin SI, Wu CC, Chiang SF, Su CL. 2006. Regulation of phosphate homeostasis by microRNA in Arabidopsis. Plant Cell 18, 412–421.

Fransz P, Linc G, Lee CR, Aflitos SA, Lasky JR, Toomajian C, Ali H, Peters J, van Dam P, Ji X, Kuzak M, Gerats T, Schubert I, Schneeberger K, Colot V, Martienssen R, Koornneef M, Nordborg M, Juenger TE, de Jong H, Schranz ME. 2016. Molecular, genetic and evolutionary analysis of a paracentric inversion in Arabidopsis thaliana. Plant J 88, 159–178.

Gong Z, Dong CH, Lee H, Zhu J, Xiong L, Gong D, Stevenson B, Zhu JK. 2005. A DEAD box RNA helicase is essential for mRNA export and important for development and stress responses in Arabidopsis. Plant Cell 17, 256–267.

Grimm DG, Roqueiro D, Salome PA, Kleeberger S, Greshake B, Zhu W, Liu C, Lippert C, Stegle O, Scholkopf B, Weigel D, Borgwardt KM. 2017. easyGWAS: A Cloud-Based Platform for Comparing the Results of Genome-Wide Association Studies. Plant Cell 29, 5–19.

Harper AL, Trick M, Higgins J, Fraser F, Clissold L, Wells R, Hattori C, Werner P, Bancroft I. 2012. Associative transcriptomics of traits in the polyploid crop species Brassica napus. Nat Biotechnol 30, 798–802.

Havlickova L, He Z, Wang L, Langer S, Harper AL, Kaur H, Broadley MR, Gegas V, Bancroft I. 2018. Validation of an updated Associative Transcriptomics platform for the polyploid crop species Brassica napus by dissection of the genetic architecture of erucic acid and tocopherol isoform variation in seeds. Plant J 93, 181–192.

Herr AJ, Molnar A, Jones A, Baulcombe DC. 2006. Defective RNA processing enhances RNA silencing and influences flowering of Arabidopsis. Proc Natl Acad Sci U S A 103, 14994–15001.

Huo H, Wei S, Bradford KJ. 2016. DELAY OF GERMINATION1 (DOG1) regulates both seed dormancy and flowering time through microRNA pathways. Proc Natl Acad Sci U S A 113, E2199–2206.

Johanson U, West J, Lister C, Michaels S, Amasino R, Dean C. 2000. Molecular Analysis of FRIGIDA, a Major Determinant of Natural Variation in Arabidopsis Flowering Time. Science 290, 344–347.

Kang HM, Zaitlen NA, Wade CM, Kirby A, Heckerman D, Daly MJ, Eskin E. 2008. Efficient control of population structure in model organism association mapping. Genetics 178, 1709–1723.

Kawakatsu T, Huang SC, Jupe F, Sasaki E, Schmitz RJ, Urich MA, Castanon R, Nery JR, Barragan C, He Y, Chen H, Dubin M, Lee CR, Wang C, Bemm F, Becker C, O’Neil R, O’Malley RC, Quarless DX, Genomes C, Schork NJ, Weigel D, Nordborg M, Ecker JR. 2016. Epigenomic Diversity in a Global Collection of Arabidopsis thaliana Accessions. Cell 166, 492–505.

Kim D-H. 2020. Current understanding of flowering pathways in plants: focusing on the vernalization pathway in Arabidopsis and several vegetable crop plants. Horticulture, Environment, and Biotechnology 61, 209–227.

Kim D, Langmead B, Salzberg SL. 2015. HISAT: a fast spliced aligner with low memory requirements. Nat Methods 12, 357–360.

Kim S, Plagnol V, Hu TT, Toomajian C, Clark RM, Ossowski S, Ecker JR, Weigel D, Nordborg M. 2007. Recombination and linkage disequilibrium in Arabidopsis thaliana. Nat. Genet. 39, 1151–1155.

Koornneef M, Blankestijn-de Vries H, Hanhart C, Soppe W, Peeters T. 1994. The phenotype of some late-flowering mutants is enhanced by a locus on chromosome 5 that is not effective in the Landsberg erecta wild-type. Plant J. 6, 911–919.

Kovaka S, Zimin AV, Pertea GM, Razaghi R, Salzberg SL, Pertea M. 2019. Transcriptome assembly from long-read RNA-seq alignments with StringTie2. Genome Biol 20, 278.

Kremling KAG, Diepenbrock CH, Gore MA, Buckler ES, Bandillo NB. 2019. Transcriptome-Wide Association Supplements Genome-Wide Association in Zea mays. G3 (Bethesda) 9, 3023–3033.

Lan Y, Sun R, Ouyang J, Ding W, Kim MJ, Wu J, Li Y, Shi T. 2021. AtMAD: Arabidopsis thaliana multi-omics association database. Nucleic Acids Res 49, D1445–D1451.

Li D, Liu Q, Schnable PS. 2021. TWAS results are complementary to and less affected by linkage disequilibrium than GWAS. Plant Physiol.

Li P, Filiault D, Box MS, Kerdaffrec E, van Oosterhout C, Wilczek AM, Schmitt J, McMullan M, Bergelson J, Nordborg M, Dean C. 2014. Multiple FLC haplotypes defined by independent cis-regulatory variation underpin life history diversity in Arabidopsis thaliana. Genes Dev 28, 1635–1640.

Li Z, Wang P, You C, Yu J, Zhang X, Yan F, Ye Z, Shen C, Li B, Guo K, Liu N, Thyssen GN, Fang DD, Lindsey K, Zhang X, Wang M, Tu L. 2020. Combined GWAS and eQTL analysis uncovers a genetic regulatory network orchestrating the initiation of secondary cell wall development in cotton. New Phytol.

Liang Y, Liu Q, Wang X, Huang C, Xu G, Hey S, Lin HY, Li C, Xu D, Wu L, Wang C, Wu W, Xia J, Han X, Lu S, Lai J, Song W, Schnable PS, Tian F. 2019. ZmMADS69 functions as a flowering activator through the ZmRap2.7-ZCN8 regulatory module and contributes to maize flowering time adaptation. New Phytol 221, 2335–2347.

Lipka AE, Tian F, Wang Q, Peiffer J, Li M, Bradbury PJ, Gore MA, Buckler ES, Zhang Z. 2012. GAPIT: genome association and prediction integrated tool. Bioinformatics 28, 2397–2399.

Ma Y, Min L, Wang J, Li Y, Wu Y, Hu Q, Ding Y, Wang M, Liang Y, Gong Z, Xie S, Su X, Wang C, Zhao Y, Fang Q, Li Y, Chi H, Chen M, Khan AH, Lindsey K, Zhu L, Li X, Zhang X. 2021. A combination of genome-wide and transcriptome-wide association studies reveals genetic elements leading to male sterility during high temperature stress in cotton. New Phytol 231, 165–181.

Mahrez W, Shin J, Munoz-Viana R, Figueiredo DD, Trejo-Arellano MS, Exner V, Siretskiy A, Gruissem W, Kohler C, Hennig L. 2016. BRR2a Affects Flowering Time via FLC Splicing. PLoS Genet 12, e1005924.

Michaels SD, Amasino RM. 1999. FLOWERING LOCUS C Encodes a Novel MADS Domain Protein That Acts as a Repressor of Flowering. The Plant Cell 11, 949–956.

Michaels SD, Amasino RM. 2001. Loss of FLOWERING LOCUS C Activity Eliminates the Late-Flowering Phenotype of FRIGIDA and Autonomous Pathway Mutations but Not Responsiveness to Vernalization. The Plant Cell 13, 935–941.

Obayashi T, Aoki Y, Tadaka S, Kagaya Y, Kinoshita K. 2018. ATTED-II in 2018: A Plant Coexpression Database Based on Investigation of the Statistical Property of the Mutual Rank Index. Plant Cell Physiol 59, e3.

Schmalenbach I, Zhang L, Ryngajllo M, Jiménez-Gómez JM. 2014. Functional analysis of the Landsberg erecta allele of FRIGIDA. BMC Plant Biology 14.

Seren Ü. 2018. GWA-Portal: Genome-Wide Association Studies Made Easy. In: Ristova D, Barbez E, eds. Root Development: Methods and Protocols. New York, NY: Springer New York, 303–319.

Seren Ü, Vilhjalmsson BJ, Horton MW, Meng D, Forai P, Huang YS, Long Q, Segura V, Nordborg M. 2012. GWAPP: a web application for genome-wide association mapping in Arabidopsis. Plant Cell 24, 4793–4805.

Sheldon CC, Rouse DT, Finnegan EJ, Peacock WJ, Dennis ES. 2000. The molecular basis of vernalization: the central role of FLOWERING LOCUS C (FLC). Proc Natl Acad Sci U S A 97, 3753–3758.

Stinchcombe JR, Weinig C, Ungerer M, Olsen KM, Mays C, Halldorsdottir SS, Purugganan MD, Schmitt J. 2004. A latitudinal cline in flowering time in Arabidopsis thaliana modulated by the flowering time gene FRIGIDA. Proc Natl Acad Sci U S A 101, 4712–4717.

Strauch RC, Svedin E, Dilkes B, Chapple C, Li X. 2015. Discovery of a novel amino acid racemase through exploration of natural variation in Arabidopsis thaliana. Proc Natl Acad Sci U S A 112, 11726–11731.

Tang S, Zhao H, Lu S, Yu L, Zhang G, Zhang Y, Yang QY, Zhou Y, Wang X, Ma W, Xie W, Guo L. 2020. Genome- and transcriptome-wide association studies provide insights into the genetic basis of natural variation of seed oil content in Brassica napus. Mol Plant 14, 470–487.

Tenaillon MI, Sawkins MC, Long AD, Gaut RL, Doebley JF, Gaut BS. 2001. Patterns of DNA sequence polymorphism along chromosome 1 of maize (Zea mays ssp. mays L.). Proc Natl Acad Sci U S A 98, 9161–9166.

The 1001 Genomes Consortium. 2016. 1,135 Genomes Reveal the Global Pattern of Polymorphism in Arabidopsis thaliana. Cell 166, 481–491.

traLin HY, Liu Q, Li X, Yang J, Liu S, Huang Y, Scanlon MJ, Nettleton D, Schnable PS. 2017. Substantial contribution of genetic variation in the expression of transcription factors to phenotypic variation revealed by eRD-GWAS. Genome Biol 18, 192.

Vasseur F, Bontpart T, Dauzat M, Granier C, Vile D. 2014. Multivariate genetic analysis of plant responses to water deficit and high temperature revealed contrasting adaptive strategies. J Exp Bot 65, 6457–6469.

Vasseur F, Exposito-Alonso M, Ayala-Garay OJ, Wang G, Enquist BJ, Vile D, Violle C, Weigel D. 2018. Adaptive diversification of growth allometry in the plant Arabidopsis thaliana Proc Natl Acad Sci U S A 115, 3416–3421.

Zhang L, Jiménez-Gómez JM. 2020. Functional analysis of FRIGIDA using naturally occurring variation in Arabidopsis thaliana. Plant J 103, 154–165.

Zheng Z, Hey S, Jubery T, Liu H, Yang Y, Coffey L, Miao C, Sigmon B, Schnable JC, Hochholdinger F, Ganapathysubramanian B, Schnable PS. 2020. Shared Genetic Control of Root System Architecture between Zea mays and Sorghum bicolor. Plant Physiol 182, 977–991.

Zhou Z, Jiang Y, Wang Z, Gou Z, Lyu J, Li W, Yu Y, Shu L, Zhao Y, Ma Y, Fang C, Shen Y, Liu T, Li C, Li Q, Wu M, Wang M, Wu Y, Dong Y, Wan W, Wang X, Ding Z, Gao Y, Xiang H, Zhu B, Lee SH, Wang W, Tian Z. 2015. Resequencing 302 wild and cultivated accessions identifies genes related to domestication and improvement in soybean. Nat Biotechnol 33, 408–414.

